# The Hypno-PC: Uncovering Sleep Dynamics through Principal Component Analysis and Hidden Markov Modeling of Electrophysiological Signals

**DOI:** 10.1101/2025.01.02.631039

**Authors:** Miriam Guendelman, Oren Shriki

## Abstract

The conventional approach to sleep analysis relies on pre-defined, visually scored stages derived from electrophysiological signals. This manual method demands substantial effort and is influenced by subjective assessments, implicitly assuming that these categories accurately reflect underlying biological processes. Recent advancements indicate that low-dimensional representations of complex brain activity can provide objective means of identifying brain states. These approaches can potentially uncover inherent patterns within sleep, offering valuable insights into its organization.

In this study, we applied Principal Component Analysis (PCA) to spectral features extracted from high-density EEG, EOG, EMG, and ECG recorded overnight at both 30– and 4-second resolutions. Notably, the first principal component—the “Hypno-PC”—strongly aligns with the hypnogram at both time scales. Subsequently, we employed a Gaussian Hidden Markov Model (GHMM) to delineate discrete states in the PCA-transformed data and to quantify their temporal dynamics. Using minimal supervision (less than 0.5% of the data labeled) and a cross-subject approach, the model achieved alignment with standard sleep labels comparable to the typical inter-rater agreement. Finally, independent component analysis (ICA) was applied to the PCA space, decomposing it into an independent set of components that potentially represent distinct physiological processes.

The integrated use of PCA, GHMM, and ICA provides a reproducible and scalable methodology that aligns with traditional sleep staging, while offering a more flexible and comprehensive perspective on sleep states. Our findings indicate that these data-driven, unsupervised methods effectively uncover the intrinsic dynamics of sleep, advancing automated sleep analysis and enhancing our understanding of sleep organization.

**Statement of Significance:** This study introduces a data-driven framework for sleep analysis designed to objectively identify brain states using low-dimensional representations of electrophysiological signals during sleep. By integrating a suite of unsupervised learning techniques, our methodology offers an alternative to subjective manual scoring, potentially enhancing both efficiency and reproducibility. In addition to aligning with traditional sleep staging, this approach uncovers subtle sleep dynamics across multiple time scales, enabling the discovery of patterns that conventional methods might overlook. These advancements hold promise for automated sleep monitoring and the study of sleep disorders, potentially improving diagnostic accuracy and facilitating large-scale sleep research. By addressing current limitations in sleep analysis techniques, this framework lays the groundwork for more elaborate and scalable assessments.

## Introduction

Understanding the complex dynamics of brain activity is a fundamental goal in neuroscience, with significant implications for understanding brain dynamics and diagnosing and treating neurological disorders. Neuroimaging tools such as functional magnetic resonance imaging (fMRI)^1^ and electroencephalography (EEG)^2^ have been instrumental in unraveling brain dynamics—particularly in the study of sleep stages.^3^

Sleep research focuses on five stages: wake, non-rapid eye movement (NREM) 1-3, and rapid eye movement (REM). The American Academy of Sleep Medicine (AASM) classifies these stages based on polysomnographic (PSG) recordings, which include EEG, electrooculography (EOG), and electromyography (EMG).^4^ Each stage is associated with characteristic oscillatory patterns in the EEG: posterior alpha activity marks quiet wakefulness; frontal theta dominates in N1 and REM; spindle activity is a hallmark of N2; and slow waves and delta waves define N3, or slow-wave sleep (SWS). Meanwhile, EOG and EMG play key roles in distinguishing REM sleep from wakefulness. Sleep stages are strongly tied to autonomic balance; thus, heart rate and variability measures from electrocardiography (ECG) may provide essential information.^5^

Clinical polysomnography (PSG) typically employs a limited number of EEG electrodes for practical reasons, such as ease of placement and streamlined visual analysis. However, this approach limits the information captured about the spatial distribution of electrophysiological activity. High-density EEG has become a valuable tool for studying sleep architecture in greater detail.^6–10^ Nevertheless, representing such complex multidimensional data in a clear and interpretable way remains a significant challenge.^11^

Recent work utilizing whole-night fMRI has revealed the intricate dynamics of sleep, identifying multiple distinct states and highlighting the complexity of sleep architecture.^12^ From a different avenue, advancements in data analysis have demonstrated that complex brain dynamics can be robustly embedded into low-dimensional spaces, revealing integrated brain dynamics^13^. These tools offer powerful insights into various brain states during sleep. However, methods often rely on non-linear transformations, making it non-trivial to interpret and map back to the source signals. This hinders a direct understanding of the mechanisms driving these representations and limits the relation of low-dimensional representation with clinical conventions and basic research findings.

Principal Component Analysis (PCA)^14^ offers a linear, unsupervised dimensionality reduction technique that can identify the principal axes of variation in high-dimensional data. The linearity of PCA enables straightforward interpretation of the directions of major variance axes in the high-dimensional feature space. As an unsupervised method, PCA is free from the subjective biases that are associated with manual labeling. Demonstrating convergence between PCA-derived representations and standard sleep classification can validate the robustness of both approaches. Such convergence can enhance our understanding of sleep dynamics and provide a complementary, quantitative perspective for evaluating sleep.

Rather than viewing sleep as a sequence of discrete, isolated stages, it is more accurate to describe it as a continuous progression of physiological states that evolve throughout the night, as reflected in dynamic changes in neuronal oscillations. A good starting point for characterizing this progression is to examine the spectral profile of the EEG signal and then use dimensionality reduction to identify patterns of temporal co-variation within that profile. Integrating EMG, EOG, and ECG signals—which capture muscular, ocular, and cardiac activity, respectively—can further reveal a broad range of physiological changes and help provide a more holistic view of the sleep process. These additional modalities also facilitate the alignment of PCA-derived representations with established sleep-stage classifications, bridging the gap between this data-driven approach and conventional frameworks.

We applied a Gaussian Hidden Markov Model (GHMM)^15^ to the PCA-transformed data to capture temporal dynamics and classify the data into discrete states. The GHMM divides the data into a predefined set of states and determines the likelihood of each sample belonging to a particular state by considering both the observed signal (emission probability) and the probability of transitioning from one state to another (transition probability) based on the previous state. By simultaneously considering the observed data at a particular time and its temporal context, the GHMM provides a more nuanced depiction of how signals evolve over time.

This process resembles the approach taken by human scorers during sleep staging; however, unlike a human rater, the model explicitly provides the emission and transition probabilities for each state. This detailed information offers valuable insights into how each state is represented within the feature space, the typical state sequences observed during sleep, and the stability of each state. These properties could enhance the reproducibility of state classification and offer complementary information about stage stability, which may be particularly relevant in various clinical conditions.^16^

PCA assumes that variation directions must be orthogonal, yet independent processes in brain activity may not fully adhere to this assumption. Moreover, orthogonality in the feature space may not accurately capture the independent processes that occur throughout the night. To address this, the PC representation can be refined by applying a blind source separation approach to identify independent processes. For this task, Independent Component Analysis (ICA)^17^ is a suitable candidate, as it maintains linearity while also addressing this limitation of PCA. ICA decomposition can potentially isolate processes originating from distinct sources, further refining the PCA representation and reflecting underlying physiological processes.^18^ By applying ICA to the PCA-transformed data, we sought to pinpoint independent processes captured in the low-dimensional representation and examine the temporal and spatial patterns of these components.^19^

In this study, we analyzed a high-density EEG dataset of 29 healthy subjects to derive a cross-subject representation of signal co-variation during sleep. First, we applied PCA to reduce the dimensionality of the data. The first principal component closely matches the human-labeled hypnogram, reflecting major sleep stages. Next, we employed a GHMM on the PCA-transformed data at 30-second and 4-second resolutions to segment sleep into discrete states.

We used a minimally supervised approach to label these states that assigns a sleep stage to each hidden state based on the “most typical” sample (i.e., requiring only one labeled example per hidden state per training subject). Quantifying the agreement rate with the manual sleep labels, we found that the labels assigned by the model fall within the range of the reported inter-rater agreement, highlighting its robust alignment with the current sleep scoring approach. We then refined the low-dimensional representation through ICA, enabling us to isolate independent components within the PCA space and gain further insights into underlying physiological processes.

By combining the linearity and interpretability of PCA and ICA with the capacity of the GHMM for temporal modeling, we constructed a concise yet informative representation of sleep states. This framework captures the temporal dynamics of brain oscillations, along with eye, muscle, and heart activity, aligning with existing sleep analysis methodologies while providing a more precise and automated avenue for investigating both the temporal and spatial characteristics of sleep.

## Methods

### Data

A rich, high-quality dataset that captures a broad range of physiological signals during sleep. By using a well-characterized and publicly available dataset, our analyses can be more easily replicated and validated. Thus, we used the ANPHY-Sleep dataset,^6^ a publicly available resource containing polysomnographic (PSG) recordings of 29 healthy, normal sleepers. Each recording includes high-density EEG with 83 channels, two bipolar EOG channels, three bipolar EMG channels (legs and chin), and one bipolar ECG channel. The data were scored according to AASM guidelines,^4^ providing a reference hypnogram for each subject. Subject demographics such as age and gender, along with overnight sleep measures, are documented in Wei et al. 2024.^6^

### Pre-processing

Current pre-processing aims to clean the raw signals and remove non-physiological noise sources. This step provides stable inputs for subsequent feature extraction and prevents systematic artifacts from influencing the analysis. The pre-processing was performed using custom code with MNE-Python^20^ functions. All signals were initially band-pass filtered, with the frequency bands adjusted to capture information that was relevant for each signal set. The filters applied included EEG: 0.5–40 Hz, EOG: 0.3–15 Hz, EMG: 10–50 Hz, and ECG: 0.5–20 Hz.

We applied an average reference on the EEG channels. For EOG, we added a derived channel by referencing left EOG to right EOG to emphasize horizontal eye movements. Bipolar referencing was used for EMG and ECG. Signals were segmented into contiguous, non-overlapping epochs of 30 seconds and 4 seconds. These epochs form the fundamental time windows for feature extraction.

### Feature Extraction

Feature extraction converts the raw signals into numerical summaries that reflect the physiological characteristics of the signal. This step distills large volumes of data into meaningful metrics representing brain rhythms, muscle activity, eye movements, and cardiac dynamics.

For each epoch, we computed the following:

**EEG Features:** We used the Welch method to estimate the power spectral density (PSD) at 0.5– 40 Hz. Relative spectral power was extracted for low delta (0.5–1.5 Hz), high delta (1–4 Hz), theta (4–8 Hz), alpha (8–12 Hz), low sigma (10–13 Hz), high sigma (12–16 Hz), beta (15–25 Hz), and gamma (25–40 Hz) bands. The aperiodic slope and intercept (from linear fits on log-log PSD), spectral entropy, and total power were also calculated using log-transformed spectral power.

**EMG Features:** PSD between 10 and 50 Hz was computed, and total power in this range represented muscle activity.

**EOG Features:** Total spectral power between 0.3–2 Hz served as an index of eye movements.

**ECG Features:** R-peaks were detected using the SciPy “find_peaks” function,^21^ yielding R-R intervals. Heart rate (HR) and heart rate variability (HRV) metrics (SDNN and RMSSD) were computed.^22^ It is important to note that HRV measures generally provide unstable information at such short time scales. However, unlike previous studies that considered a single estimate,^22^ our approach focuses on relative changes rather than on absolute HRV values. By doing so, we utilize this instability as a feature rather than viewing it as a limitation, offering a novel perspective on HRV dynamics.

### Feature Cleaning and Standardization

This step ensures that all extracted features are on a comparable scale and that missing or extreme values do not bias downstream analyses. It stabilizes the dataset in order to apply dimensionality reduction techniques and modeling.

Within each subject, features were z-scored by subtracting the mean and dividing by the standard deviation. Missing values occurred only for the ECG measure at the 4-second resolution and in less than <0.2% of the data in 12 subjects. Most missing values were found in subject EPCTL29, where RMSSD could not be calculated for 430 segments, likely due to a low heart rate, as at least three detected peaks are required for this calculation. To maintain temporal sampling consistency, missing values were linearly interpolated. To reduce the impact of outliers on the PCA features, values were clipped to the range of [-5, 5]. Across features, the mean portion of clipped values was lower than 1% of the segments, and no trend was observed in individual features.

### Principal Component Analysis (PCA)

PCA reduces the complexity of the dataset by identifying orthogonal directions (principal components) of maximal variance. This step condenses a large number of features into a handful of variables, making the predominant patterns in the data easier to visualize and analyze.

PCA was applied to the standardized feature matrix. Components explaining at least 2% of the total variance were retained, resulting in five principal components (PCs). These PCs were plotted over time and compared to the hypnogram to interpret their relation to known sleep stages.

### Visualizing the Component Filters

This visualization aims to make the principal and independent components more interpretable by showing how each feature contributes to a component linking the abstract PCs back to meaningful physiological patterns. For EEG, spatial topography was generated by projecting the component weights of each feature onto a head map according to the electrode layout. Non-EEG features (EOG, EMG, ECG) were represented in bar plots. The color and scale of these maps highlight which features heavily influence each component, guiding physiological interpretation.

### Statistical Analysis

This analysis quantified the relationship between each component and known sleep stages. It tested whether the PCs (or later ICs) reliably differ across these stages. We fitted a mixed-effects linear model for each component using Statsmodels^23^ with the following formula:

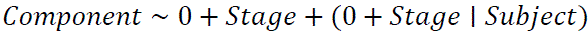

The model estimates the direction and strength of the association between each component and sleep stages while accounting for individual differences and repeated samples and estimating both the individual and overall slopes. We extracted the coefficients, p-values, standard errors, and 95% confidence intervals and visualized significance using conventional markers and plotting patient-specific slopes to assess cross-subject consistency.

### Gaussian Hidden Markov Model (GHMM)

While traditional sleep staging relies on pre-defined categories, the goal here was to discover underlying states directly from the data. The GHMM treats the transformed data as observations generated by a sequence of hidden states, each with its own emission (observation) distribution. By fitting a GHMM, we aimed to identify discrete states that reflect inherent patterns in the data—potentially corresponding to, but not dictated by, conventional sleep stages. This method allowed us to model the distribution of observations within each state and the temporal transitions between them, providing a time-resolved, data-driven perspective on sleep architecture.

We applied the GHMM to the PCA-transformed data at both 30-second and 4-second resolution epochs, allowing us to investigate the dependence of state segmentation on the temporal scale of analysis. We implemented a leave-one-subject-out cross-validation scheme. For each run, the data from 28 subjects served as training data, and the remaining subject was held out as test data. This process was repeated 29 times, ensuring that each subject was used as the test set exactly once.

GHMMs were trained with a range of possible state counts (2 to 15). For each state count, we evaluated the Bayesian information criterion (BIC) and Akaike information criterion (AIC) for the held-out subjects. The optimal number of states was chosen where increasing the number of states no longer improved either the median BIC or AIC, balancing model complexity and fit quality.

### Minimal Supervision for Label Assignment

We used a minimally supervised approach to link hidden states to known sleep stages. For each candidate state in the training set, we identified the segment within that state with the highest posterior probability—representing the most “typical” example of that state. We then examined the sleep stage labels of these most typical segments across training subjects. The sleep stage that appeared most frequently for each state was assigned as its label. This procedure uses fewer than 0.5% of the labeled segments in the data, significantly reducing manual labeling overhead. This ensures that states are primarily defined by the data, with only a minimal reference to known stages.

### Agreement Evaluation with Manual Labels

After assigning labels to the hidden states, we evaluated how well the GHMM-assigned labels aligned with the manual sleep labels in each model configuration (ranging from 2 to 15 states). Specifically, we computed both accuracy and Cohen’s kappa to quantify agreement with conventional sleep scoring.

### Comparing Different Modeling Approaches

After determining the optimal number of states, we explored three strategies for training the hidden state models. First, we used our original cross-validation approach, training each model on a subset of subjects and validating it on the remaining subjects. Second, we trained a single, unified model using data from all subjects. Finally, we adopted a personalized strategy, in which we started with the unified model and then fine-tuned it on the data of each subject prior to evaluation. We assessed the performance of these three approaches across all subjects, focusing on both the median performance and the variability observed among individuals.

At a group level, we compared the GHMM-derived overnight measures—total sleep time (TST), sleep onset latency (SOL), wake after sleep onset (WASO), sleep efficiency (SE), and time spent in each stage—with the corresponding measures obtained from manual hypnograms. We then computed inter-class correlation coefficients (ICCs),^24^ and p-values to evaluate how consistently the GHMM-based measures aligned with conventional sleep scoring.

To better understand the relationship between GHMM states and conventional sleep stages, we assessed the proportion of each standard sleep stage within each hidden state, the spatial distribution of these states in the principal component space, their occurrence throughout the night, and the transition probabilities among them. We then compared these GHMM-derived properties with the corresponding features derived from conventional sleep stages.

### Independent Component Analysis (ICA)

ICA was applied to further decompose the principal components (PCs) into statistically independent processes, helping to isolate distinct physiological generators that might be mixed within the PCs. To ensure robust separation and identifiability, each PC was re-scaled before ICA training, and the model was fitted on the five PCs using Python-Picard, which implements an extended infomax algorithm to separate both sub– and super-Gaussian signals.^25^ After fitting, the ICA unmixing transformation was applied without unit scaling to preserve the original variance ratios. We then analyzed the resulting independent component (IC) time series in the same manner as the PCs, employing topographic maps and mixed-effects modeling as described in previous sections.

## Results

Applying PCA to the extracted features (EEG, EOG, EMG, and ECG) at the 30-second timescale yielded five PCs, each explaining at least 2% of the variance. The first PC (PC1) accounted for 41.8% of the variance, suggesting that it captures a dominant pattern in the electrophysiological signals. Strikingly, when examining this PC over time in individual subjects, we found that it closely resembled the manually scored hypnogram (Figure 1A-B). This strong alignment was observed across subjects, motivating us to refer to PC1 as the “Hypno-PC.”

**Figure 1.**
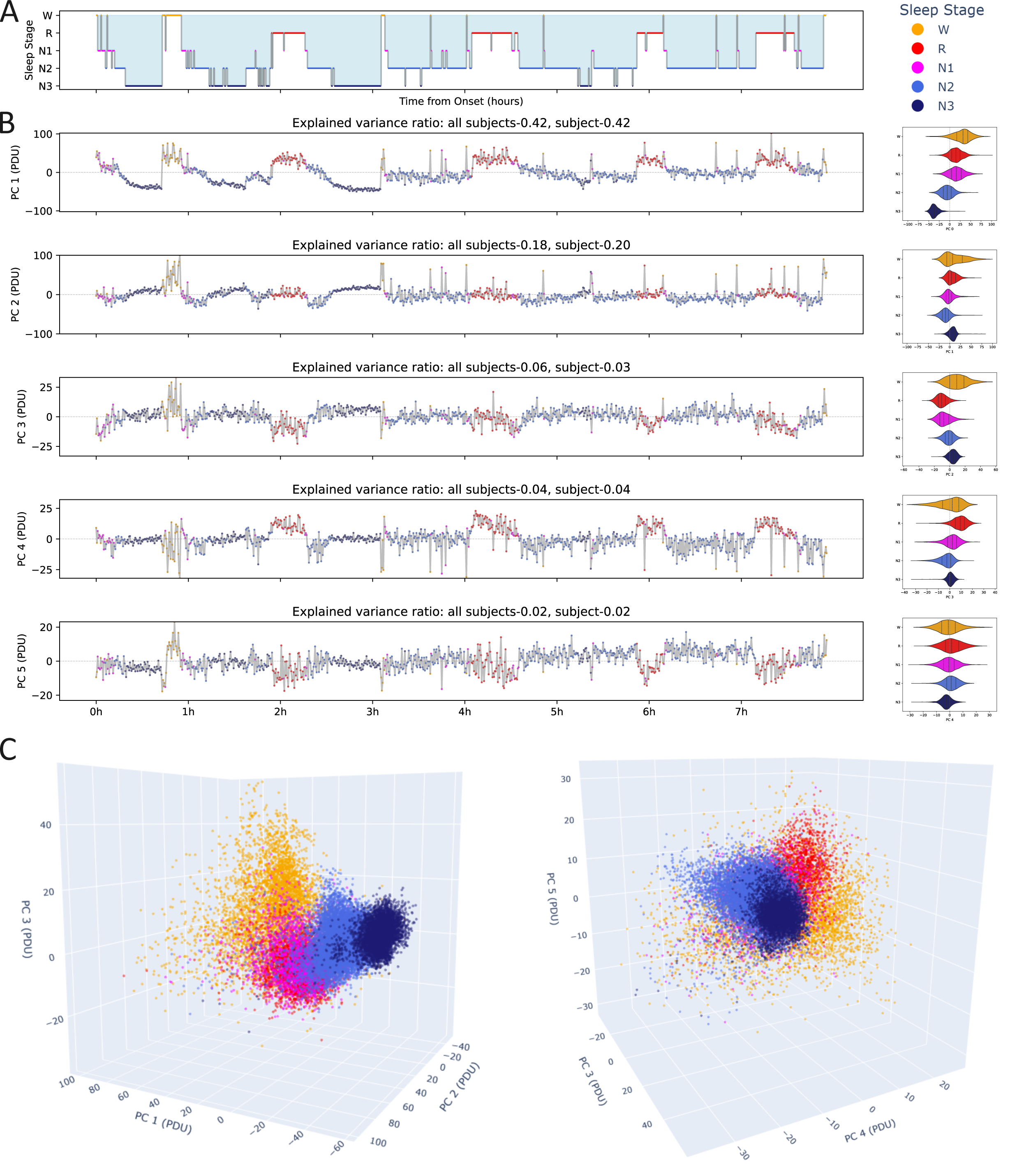
– Principal Component Timeline and Sleep Stages This figure illustrates the relationship between principal components derived from the 30-second segment decomposition of sleep data. A similar plot for the 4-second segments is provided in Figure S1. A. The manually scored hypnogram for a representative subject (EPCTL01). B. Left Panel: Time series of Principal Components 1 through 5 (PC1–PC5) for the representative subject, highlighting the strong similarity between PC1 (termed the “Hypno-PC”) and the hypnogram Right Panel: Cross-subject distributions of component values for each sleep stage, demonstrating that PC1 systematically distinguishes sleep stages across individuals. Additionally, an example time series and the overall value distribution by stage are shown for each component. C. 3D Representation: Cross-subject 3D plots of PC 1–3 and 3–5, illustrating how sleep stages occupy distinct regions within the principal component space. Notably, there is some overlap between N1 and REM sleep, as evidenced in panels A and B. Abbreviations: PC: Principal component, Hypno-PC: Principal component 1, aligned with the hypnogram, N1–3: Non-rapid eye movement 1–3, REM: Rapid eye movement sleep, W: Wake Note: All principal components are derived from spectral features of high-density electroencephalography (EEG), electrooculography (EOG), electromyography (EMG), and electrocardiography (ECG) recordings.

Beyond PC1, the remaining PCs together captured more nuanced and mixed variations in the data. For example, PCs 2 and 3 provided additional axes along which different sleep stages were arranged within the low-dimensional space (Figure 1C). While the wake, N2, and N3 stages occupied relatively distinct positions, the separation between N1 and REM was less clearly defined. These overlaps highlight the complexity and continuity of sleep physiology rather than discrete boundaries between stages.

At the 4-second resolution, a similar PCA structure emerged (Figure S1). Although PC1 still tracked the hypnogram-like pattern, the increased temporal resolution introduced greater within-stage variability. Additionally, certain patterns observed at 30-second epochs—for instance, a clearer REM signature in PCs 3–5—were less distinct at 4 seconds. The ground-truth labels were provided for 30-second windows and were resampled for this evaluation. Thus, the manual labels may not reflect possible changes at this timescale.

### Feature Maps and Statistical Modeling

To associate each principal component with specific electrophysiological processes, we created “filter maps” that depict the spatial distribution of weights for each feature. Additionally, we assessed the statistical relationship between each PC and the sleep stages using mixed-effects models. For PC1, which explained 41.8% of the variance and closely mirrors the sleep stage values in the hypnogram, positive values were associated with higher-frequency power bands, primarily beta power. Conversely, negative values corresponded to lower-frequency power bands, mainly log-delta power. This relationship was further supported by positive weights on the spectral slope, indicating that flatter (less negative) spectral slopes are linked to higher PC1 values (see Figure 2 and its caption for details on additional PCs).

**Figure 2.**
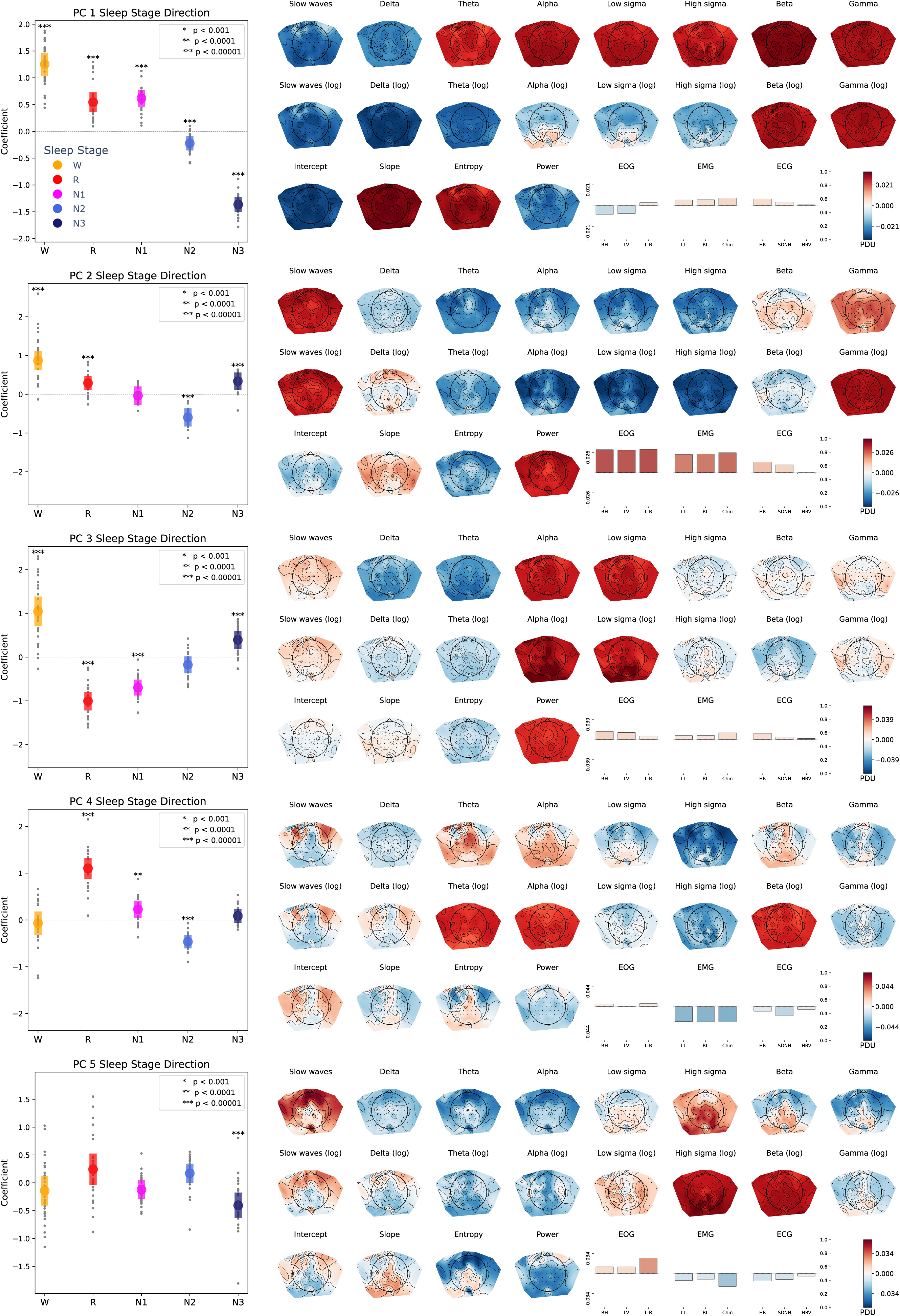
– Principal Component Filter Maps This figure depicts the principal component (PC) filters derived from high-density electroencephalography (EEG), electrooculography (EOG), electromyography (EMG), and electrocardiography (ECG) recordings during sleep. Each PC functions as a linear filter applied to the high-dimensional feature space, highlighting the contributions of different frequency bands, electrodes, and physiological measures. **PC1 (Hypno-PC):** Primarily differentiates wakefulness and slow-wave sleep. Positive (red) values are associated with higher-frequency EEG activity, which is characteristic of wakefulness and rapid eye movement (REM) sleep. In contrast, negative (blue) values correspond to increased lower-frequency activity, indicating deeper non-REM (NREM) sleep. **PC2:** Negative values emphasized N2 sleep, with features linked to the theta-to-sigma range. Positive values drew elements from N3 (slow waves), REM (eye movements), and wakefulness (muscle movement and high-frequency activity) states, reflecting a mixture of multiple processes. **PC3:** Strongly related to a wake-REM contrast, with positive values associated with posterior alpha (wake) and negative values related to REM and N1, and associated with an increase in theta and delta values. This component does not strongly relate to eye movement and thus may be related to tonic REM. **PC4:** Most positive for REM sleep. In this direction, there is an increase in bilateral frontal slow waves, central theta, and posterior alpha and beta, which are patterns observed in REM. Muscle activity drives the component to negative values, the opposite of REM and a trend that aligns with REM atonia. **PC5:** Negatively correlated with N3 sleep. Positive values are more related to REM and N2 sleep, featuring horizontal movement, anterior slow waves, posterior 10-25 Hz oscillations, and an increased posterior spectral slope, particularly in the signal range. Similar patterns have been associated with dream experiences.^29^ *=p < 0.001, **=p < 0.0001, ***=p<0.0001, Abbreviations: PC: Principal component, EEG: Electroencephalography, EOG: Electrooculography, EMG: Electromyography, ECG: Electrocardiography, REM/R: Rapid eye movement sleep, NREM: Non-rapid eye movement sleep, N1-3: NREM1-3, W: wake. Note: All principal components are derived from spectral features of high-density EEG, EOG, EMG, and ECG recordings during sleep.

At the 4-second resolution, the spatial patterns and directional relationships remained broadly consistent for PCs 1–3. However, PCs 4–5 displayed less defined spatial structures and appeared more sensitive to high-frequency spatial variability at this finer temporal scale (see Figure S2 for more detail).

### Identifying Inherent States Using a Gaussian Hidden Markov Model (GHMM)

To identify inherent states in the data and to assess their temporal dynamics, we applied a GHMM to the PCA-transformed data. The GHMM framework assumes that the observed signals are generated as emissions from a series of hidden states that evolve sequentially over time. In this context, emission refers to the measured signals or data points that are produced by the underlying, unobserved (hidden) states.

We trained GHMMs with varying numbers of states (ranging from 2 to 15) and employed a leave-one-subject-out cross-validation strategy to determine the optimal state number across subjects. For the 30-second epoch, a four-state model provided the best balance between model complexity and fit quality, as indicated by both the BIC and AIC. At the finer 4-second resolution, a seven-state model emerged as optimal. These results, detailed in Figure S3, demonstrate that different temporal scales can uncover varying levels of granularity in sleep structure, highlighting the nuanced dynamics captured by the GHMM approach.

### Minimal Supervision and Agreement with Labels

To map the hidden states to conventional sleep stages, we used a minimally supervised labeling approach. For each hidden state and each subject in the training set, we identified the most representative segment—defined as the segment with the highest posterior probability of belonging to that state. We then assigned each hidden state the most frequent sleep stage label among these representative segments across all training subjects. Notably, this procedure required less than 0.5% of the labeled data.

Despite such sparse supervision, the GHMM approach achieved an inter-rater-level agreement rate within the reliability range commonly seen among expert human scorers^26^ (kappa ≈0.71 at 30-second resolution). This result shows that the GHMM effectively captured the underlying structure of the data and that this structure generally aligns with clinical sleep staging, thus providing a reliable and interpretable automated approach to estimating sleep stages.

### Modeling Approaches and Overnight Measures

To evaluate how training the approach affected the alignment of the sleep labels and GHMM states, we compared three GHMM setups:

1. Cross-Subject (Leave-One-Out) Approach: Each run was trained on 28 subjects and tested on the remaining subject, providing a robust measure of generalizability.
2. Shared Model: A single model trained on all subjects simultaneously, creating a global model of sleep structure.
3. Personalized Model: Here, the shared model was fine-tuned based on the data of each subject prior to evaluation.

At a 30-second temporal resolution, the shared model demonstrated slightly higher consistency with manual scoring than the other approaches. Interestingly, personalized models—designed to capture individual sleep nuances—not only did not improve agreement with standard labels, but they also often diminished it. This indicates that there are key elements in sleep structure that are shared across individuals that align with the conventional criteria. Refining the fit on a particular subject that highlights personal idiosyncrasies reduces the agreement with conventional criteria (Table 1, Top). Nonetheless, exploring the individual variations and the divergence from “normal” sleep architecture using personalized models may provide insights into different sleep pathologies.

**Table 1.** Agreement Metrics and Intraclass Correlation (ICC) Values for Overnight Sleep Measures across Modeling Approaches at 30 and 4 Second Resolutions.

| Metric | TST | SOL | WASO | SE | NREM 1 | NREM 2 | NREM 3 | NREM | REM | Kappa | Accuracy |
| --- | --- | --- | --- | --- | --- | --- | --- | --- | --- | --- | --- |
| 30 second 4-stage model (ICC2) |  |  |  |  |  |  |  |  |  | Mean [SD] |  |
| Cross subject | 0.954<br>*** | 0.949<br>*** | 0.874<br>*** | 0.901<br>*** | -0.058<br>ns | 0.726<br>*** | 0.466<br>* | 0.622<br>*** | 0.264<br>** | 0.701<br>[0.067] | 0.781<br>[0.059] |
| Shared model | 0.959<br>*** | 0.949<br>*** | 0.885<br>*** | 0.916<br>*** | 0<br>ns | 0.762<br>*** | 0.391<br>* | 0.633<br>*** | 0.281<br>*** | 0.710<br>[0.07] | 0.787<br>[0.054] |
| Personalized model | 0.871<br>*** | 0.981<br>*** | 0.734<br>*** | 0.751<br>*** | -0.001<br>ns | 0.407<br>* | 0.134<br>ns | 0.653<br>*** | 0.330<br>Ns | 0.662<br>[0.138] | 0.747<br>[0.120] |
| 4 second 7-stage model (ICC2) |  |  |  |  |  |  |  |  |  | Mean [SD] |  |
| Cross subject | 0.951<br>*** | 0.948<br>*** | 0.881<br>*** | 0.903<br>*** | 0.042<br>ns | 0.649<br>*** | 0.272<br>ns | 0.738<br>*** | 0.167<br>Ns | 0.669<br>[0.087] | 0.758<br>[0.066] |
| Shared model | 0.958<br>*** | 0.741<br>*** | 0.883<br>*** | 0.913<br>*** | 0.253<br>ns | 0.465<br>*** | 0.426<br>*** | 0.835<br>*** | 0.041<br>ns | 0.615<br>[0.091] | 0.709<br>[0.067] |
| Personalized model | 0.922<br>*** | 0.934<br>*** | 0.854<br>*** | 0.878<br>*** | -0.01<br>ns | 0.716<br>*** | 0.376<br>ns | 0.766<br>*** | 0.411<br>Ns | 0.627<br>[0.120] | 0.730<br>[0.097] |

Symbols: ns=not significant, \*=p < 0.01, \*\*=p < 0.001, \*\*\*=p<0.001,
Abbreviations: TST: Total sleep time, SOL: Sleep onset latency, WASO: Wake after sleep onset,
SE: Sleep efficiency, NREM: Non-rapid eye movement sleep, REM: Rapid eye movement sleep
Note: All values are presented as Mean [Standard Deviation] where applicable.

At the 4-second scale, absolute agreement was lower, which is expected given that ground-truth labels were not originally defined at this temporal resolution (Table 1, bottom). The shared model is the only one where N1 is consistently included. In Figure 5, we can see that in this model, both the N1 and REM-related stages are highly heterogeneous, with a high proportion of wake and REM-labeled segments in the state assigned to N1 and a high portion of N1 and N2 in the state assigned to REM. These heterogeneities may result in short temporal patterns (e.g., brief frontal theta or alpha waves, tonic REM phases) that are typically observed within the 30-second segments.

This heterogeneity led to underestimates of SOL and overestimates of REM time, yet each model continued to capture important features of sleep. For instance, TST, WASO, and sleep efficiency SE exhibited strong ICC values across approaches, demonstrating that the GHMM remains valuable for capturing essential aspects of overnight sleep architecture.

### Understanding Sleep and Hidden State Characteristics

To further explore how the GHMM states relate to conventional sleep stages, we examined the distribution of labels within each hidden state. We visualized the temporal evolution of these states overnight. By considering emission and transition probabilities, the GHMM provides insights into the temporal dynamics of the states throughout the night. Figure 3 illustrates temporal metrics derived from the hypnogram and highlights two illustrative examples showing the importance of stability and transition measures. Figure S4 demonstrates common temporal patterns captured from the 4-second model in these subjects.

**Figure 3.**
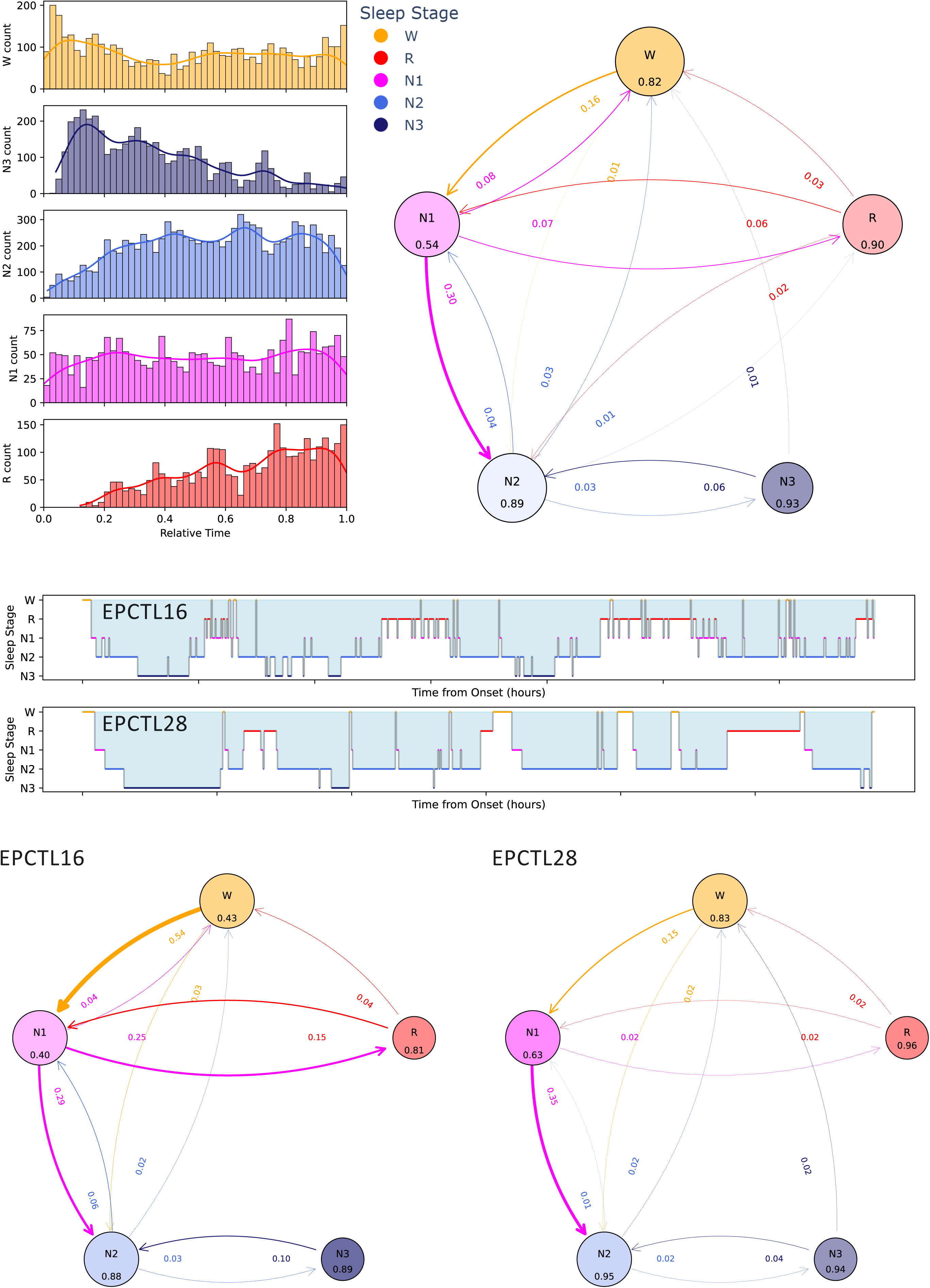
– Temporal Analysis of Sleep Stage Transitions This figure examines the temporal dynamics of sleep stages across subjects, highlighting both overall patterns and individual variations. **Top-left Panel:** Distribution of conventional sleep stages throughout the night, illustrating common patterns such as an early dominance of N3 and a later increase in N2 and REM sleep. **Top-right Panel:** Summary of stage-to-stage transitions averaged across subjects. Each circle represents a sleep stage, with its size proportional to the number of transitions into that stage. The color saturation decreases as outgoing transitions increase, providing a visual representation of the number of possible transitions and connectivity to other stages. The number within each circle represents the stage stability (self-transitions), e.g., how likely it is to remain in this stage across consecutive windows. **Bottom Panel:** A comparison of two subjects (EPCTL16 and EPCTL28) exhibited high sleep efficiency (96% and 91%, respectively). Although EPCTL28 has a shorter Total Sleep Time (TST = 321.5 minutes vs. 392.5 minutes) and slightly more Wake After Sleep Onset (WASO = 27.5 minutes vs. 13 minutes), conventional metrics alone do not fully capture the complexity of their sleep. The hypnogram for EPCTL16 indicates more fragmented sleep with frequent transitions among the REM, N1, and N2 stages. Transition probability plots reflect these observations, showing reduced state stability and greater flux between N1 and REM in EPCTL16 compared to EPCTL28. This comparison demonstrates how visualizing transitions provides additional context beyond standard sleep efficiency metrics, revealing nuances in sleep architecture and stability. Abbreviations: REM/R: Rapid eye movement sleep, NREM: Non-rapid eye movement sleep, N1–3: NREM1–3, W: Wake, TST: Total sleep time, WASO: Wake after sleep onset Note: The data presented in this figure are extracted from the manual sleep label sequence.

The characteristics of the final chosen models (four states at 30 seconds, seven states at 4 seconds) are presented in Figures 4 and 5. These figures show how these data-driven states reflect established sleep patterns while capturing subtler transitions and variations that classical staging methods might overlook^27^. Moreover, using the emission probabilities, we visualize the relationship between the PC values and the different stages.

**Figure 4.**
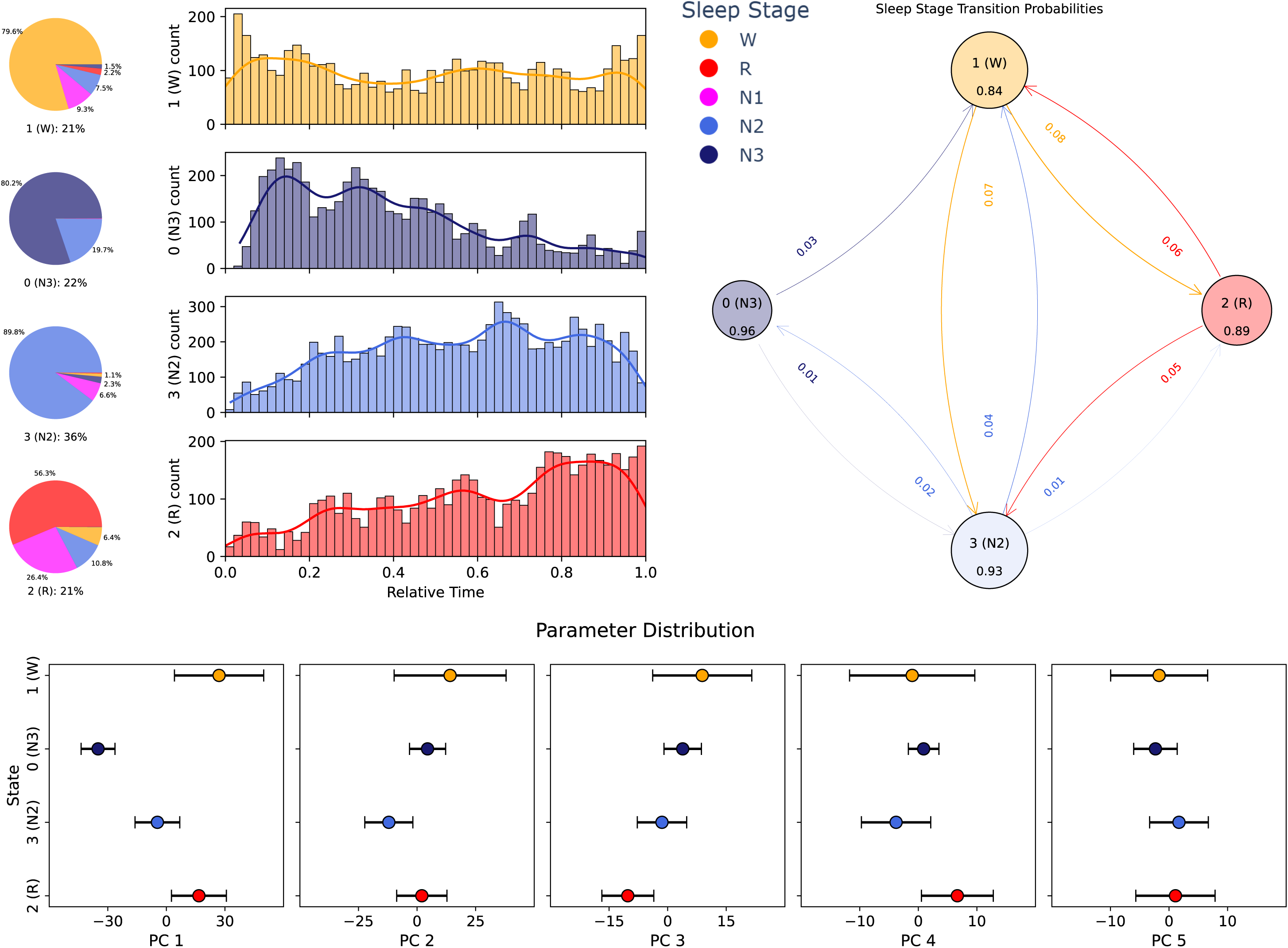
– Properties of Hidden States Identified by Gaussian Hidden Markov Model at 30-Second Resolution This figure illustrates the properties of the hidden states identified by the Gaussian hidden Markov model (GHMM) at the 30-second resolution. **Top-left panel:** The proportion of standard sleep stages within each discovered hidden state. Four states were identified: ***State 1 (W):*** Constitutes 21% of the data, with 79.6% of these segments labeled as Wake, 9.3% as N1, and 7.5% as N2. ***State 0 (N3):*** Encompasses 22% of the data, primarily composed of N3 segments (80.2%) alongside some N2 segments (19.7%). ***State 3 (N2):*** Accounts for 36% of the data, with 89.8% labeled as N2 and a minor fraction (6.6%) as N1. ***State 2 (REM):*** Includes 21% of the data. Although REM dominates at 56.3%, N1 (26.4%), N2 (10.8%), and Wake (6.4%) also appear. This mixture reflects the close physiological similarity between these stages, as recognized by AASM guidelines. An N1-related state (9.8% of the data) was not identified. Segments labeled with this state were most commonly found in the REM-like state (5.5%), and at a comparable rate in the wake (2%) and N2 (2.4%) states. **Top-middle panel:** The overnight occurrence rate of these states shows that their temporal distribution aligns well with conventional stage patterns presented in Figure 3. **Top-left panel:** The transition probabilities among the hidden states. Each circle represents a hidden state, with its size proportional to the number of transitions into that stage. The color saturation decreases as outgoing transitions increase, providing a visual representation of the number of possible transitions and connectivity to other stages. The number within each circle represents the stage stability (self-transitions), e.g., how likely it is to remain in this stage across consecutive windows. The overall stability of the hidden states was higher compared to the manual hypnogram stages; this may be due to the reduction of the impact of the N1-related transitions observed in the hypnogram (e.g., W-N1, N1-N2), suggesting better-defined states. **Lower panel:** Distribution of hidden states within the principal component (PC) space. Comparing their spatial arrangement to that presented in Figures 1 and 2 illustrates how GHMM states align with the distribution of sleep stages in the PC space. Despite some label misalignment, this visualization demonstrates that GHMM-derived states preserve key temporal and spatial characteristics of established sleep stages while providing a more stable, data-driven representation. Abbreviations: REM/R: Rapid eye movement sleep, NREM: Non-rapid eye movement sleep, N1–3: NREM1–3, W: Wake, GHMM: Gaussian hidden Markov model, PC: Principal component, AASM: American Academy of Sleep Medicine.

**Figure 5.**
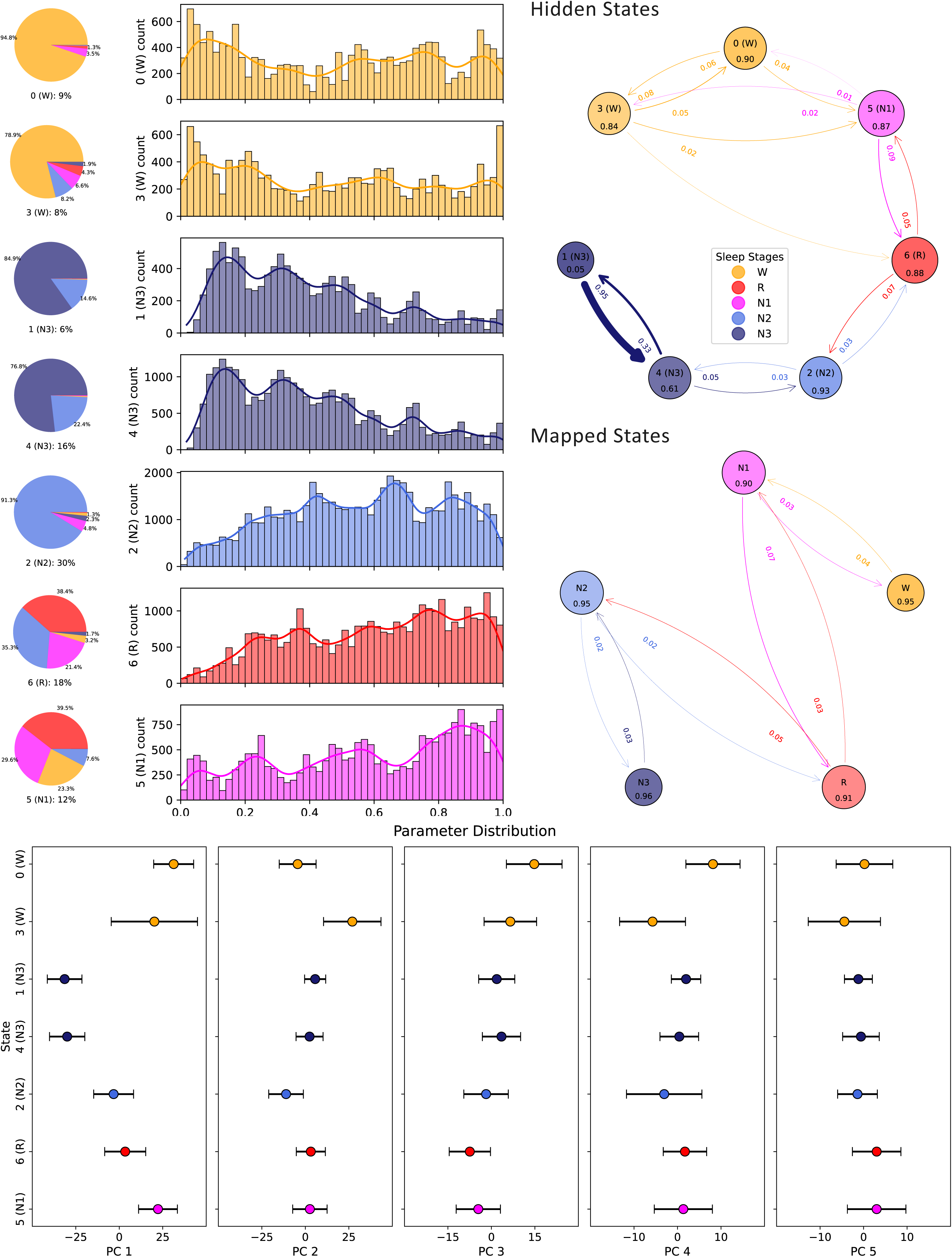
– Properties of Hidden States Identified by Gaussian Hidden Markov Model at 4-Second Resolution This figure illustrates the properties of the hidden states identified by the GHMM at the 4-second resolution. **Top-Left Panel:** The state distribution plots reveal that some hidden states align well with their conventional sleep stages (e.g., those corresponding to W, N2, and N3). In contrast, the states associated with N1 and REM are more heterogeneous, containing many segments from all stages except for N3 labeled segments. Thus, these could reflect sub-states at this timescale that transiently occur within the 30-second epochs labeled as W, R, N1, or N2. **Top-Middle Panel:** The temporal distribution of these states throughout the night shows expected patterns: wake-related states predominate at the beginning and end of the night, and N3-related states cluster in a more phasic manner early on. In contrast, N1, N2, and R-related states gradually rise as the night progresses. This pattern suggests that, even at shorter timescales, the GHMM captures temporal structures that align with accepted sleep architecture. **Top-Right Panel:** With more than one state per sleep stage, both hidden state and sleep stage dynamics are evaluated in two ways: 1. Hidden State Transitions: Transition probabilities among the hidden states are determined using the GHMM model. 2. Sleep Stage Transitions: Transition probabilities are estimated from the mapped states sequence after label assignment using the minimally supervised approach and averaged across patients. Each circle represents a hidden state (top) or a sleep stage (bottom), with its size proportional to the number of transitions into that state/stage. The color saturation decreases as outgoing transitions increase, providing a visual representation of the number of possible transitions and connectivity to other stages. The number within each circle represents the stage stability (self-transitions), reflecting the likelihood of remaining in the same stage across consecutive windows. Hidden states 1 and 4, aligned with N3, exhibit low stability with a high transition rate between these stages, suggesting strongly alternating N3 substrates. Additionally, there appears to be a clear directional flow among the hidden states, primarily forming a chain-like structure. When represented as sleep stages, we can see that the overall stage stability is high, demonstrating how integrating prior knowledge (sleep labels) and data-driven segmentation produces better-defined states. Deviations from these stable patterns can serve as potential markers of sleep stability at the individual level (see Figure S4). **Lower Panel (State Distribution in PC Space):** This panel maps the probability of each hidden state within the principal component (PC) space. Comparing the REM and N1-related states, the N1-related state aligns with higher values in PC1. It includes a significant proportion of wake segments, suggesting it acts as a transitional phase from wakefulness and reflects a higher arousal state. Similarly, the REM-related state blends N1, R, and N2 features, reflecting a shared sub-state among sleep stages. Examining the two wake states shows differences primarily in PC2 and PC4 (see Figure S2 for the component filters). One wake state (hidden state 3) correlates with increases in spectral slope, gamma, slow-wave activity, and an increase in eye/muscle movement and heart rate (positive PC2), possibly indicating a more activated form of wakefulness. In contrast, the other wake state (hidden state 0) is associated with heightened theta and alpha power, suggesting a quieter wake state. A similar pattern emerges with the two N3 states: one state (4) is relatively stable and mediates transitions to and from N2, whereas the other (1) is less stable and frequently transitions back to state 4. Although these two N3 states occupy close locations in the PC space, their distinct transition patterns hint at finer-grained differences within deep sleep at this temporal resolution. Overall, these hidden states and their transitions may represent previously described phenomena, such as cyclic alternating patterns (CAP).^37^ Abbreviations: REM: Rapid eye movement sleep, NREM: Non-rapid eye movement sleep, N1-3: NREM1-3, W: Wake, GHMM: Gaussian hidden Markov model, PC: Principal component, CAP: Cyclic alternating pattern

To further dissect the low-dimensional representation obtained through PCA, we performed an ICA. As detailed in the Supplemental Information (Figures S5 and S6), ICA unmixed overlapping physiological signals (e.g., arousals, phasic REM fluctuations) that may be conflated in PCA space. Although these ICA findings do not substantially alter the stage-level conclusions drawn from PCA and GHMM, they enhance our physiological interpretation by capturing partially independent processes.^3,28,29^ These observations reinforce the complexity of sleep-state transitions and underscore the latent dynamics that shape wakefulness, NREM, and REM episodes at both temporal and spatial scales.^27,30–33^

## Discussion

Recent advances in neuroscience have shown that complex brain activity, recorded from large populations of neurons, can often be described within low-dimensional spaces,^13^ providing an enhanced understanding of the computations performed by the neuronal population.^34^ Dimensionality reduction techniques—ranging from linear methods such as principal component analysis (PCA) and factor analysis to non-linear methods like t-SNE and variational autoencoders—have been increasingly applied to characterize brain states and transitions within those states.^13^

In the context of sleep, studies have employed these approaches across multiple modalities. For example, recently, non-linear manifold learning techniques have been applied to fMRI data to uncover subtle neural dynamics underlying sleep stages.^12,35^ PCA decomposition has successfully revealed robust patterns from EEG that parallel conventional sleep staging categories from only three EEG electrodes.^36^ These efforts exemplify how low-dimensional representations can capture essential features of sleep architecture.

In this study, we introduced a data-driven framework that integrates PCA and GHMMs to analyze the inherent structure of high-density electrophysiological sleep data. Our results demonstrate that the first principal component (PC1), or “Hypno-PC,” strongly aligns with the manually scored hypnogram, confirming that the largest axis of variance in our spectral feature space closely reflects established sleep staging criteria consistent with previous findings.^36^ This convergence is particularly notable, as it emerged from a purely unsupervised and linear decomposition, underscoring the robustness of conventional stage definitions and the capacity of data-driven approaches to recover similar structures independently.

By employing a minimally supervised GHMM trained on PCA-transformed features, we classified sleep states at both 30-second and 4-second resolutions. Remarkably, using less than 0.5% of the training data’s labels, this model achieved agreement levels with manual staging comparable to typical inter-rater reliability,^24^ reducing labeling overhead and enhancing reproducibility. Furthermore, applying ICA to the PCA space allowed us to isolate independent physiological processes that underlie the observed variance, offering richer insights into the complexity of sleep dynamics beyond the standard staging categories.

### Implications for Sleep Research

These findings complement and extend traditional manual staging approaches, which—despite their clinical ubiquity—can be time-consuming, subjective, and prone to inter-rater variability.^24,26^ Our unsupervised framework not only corroborates standard stage boundaries but also quantifies subtle within-stage variability in the spatial and temporal organization. By unveiling finer temporal structures (e.g., a 4-second scale, Figure 5) and identifying stable states or transient sub-states, our approach moves toward a more continuous and fine-grained temporal representation of sleep states. Overall, at the short analysis time scale, these transitions and micro-states captured as the hidden states may represent previously described phenomena in sleep stability analysis, such as cyclic alternating patterns (CAP).^37^

Our approach not only provides temporal flexibility but also uses high-density EEG recordings to create interpretable maps of spatial covariation of the different brain oscillations, enhancing the understanding of sleep-related processes from a spatial perspective.^7,11^ Such an understanding aligns well with the notion that sleep involves complex interactions among neural oscillations, muscle activity, and autonomic regulation.^5,2,27^ By embracing these continuous, data-driven methods, researchers can gain new perspectives on how states evolve and interact at different timescales, improving our foundational models of sleep architecture.

Our transition probability analysis further underscores the ability of data-driven methods to capture the probabilistic nature of sleep state transitions. This capability is crucial for understanding the dynamics of sleep continuity and fragmentation, which are key indicators of sleep quality and health.^16^

### Clinical Applications

From a clinical standpoint, automated, data-driven methods for characterizing sleep could facilitate routine analyses in hospital sleep laboratories and potentially extend to home monitoring setups. Indeed, it has been demonstrated that similar patterns emerge with three EEG electrodes in a different cohort.^36^ We show that these may be improved by adding the EMG, EOG, and ECG that are typically recorded in a PSG. This approach could reduce the burden of manual scoring, increase labeling consistency, and detect more subtle changes in sleep patterns that may be clinically relevant.^38^

Conditions characterized by disrupted sleep continuity or altered stage stability—such as insomnia, sleep apnea, circadian rhythm disorders, and age-related changes—may be better understood by examining data-driven state dynamics,^16,39^ particularly by leveraging transition probability graphs to assess sleep stability. Moreover, conditions such as REM sleep behavior disorder,^40^ Parkinson’s disease,^41^ autism spectrum disorder,^42^ and Alzheimer’s disease^43^ are often associated with a reduction in sleep disruptions. By quantifying the state transitions objectively, our GHMM-based framework offers diagnostic information and may help tailor therapeutic strategies aiming to stabilize particular sleep stages. In other disorders such as epilepsy,^44^ linking the way pathological events (e.g., seizures and interictal epileptiform activity) to their representation in the low dimensional space and in relation to the hidden states could pave the way for a better understanding of the interaction between these pathological manifestations and brain states.

### Methodological Considerations and Future Directions

While this study advances the application of unsupervised methods in sleep research, several limitations warrant consideration. First, we relied on pre-defined segment lengths (30-second and 4-second epochs), which may not fully capture non-stationary or scale-invariant sleep processes. Evaluating different time scales or adaptive segmentation strategies, such as change-point detection,^45^ might reveal novel states or transitions previously obscured by fixed epochs.

In this work, the small portion of missing data were handled via interpolation—a potential source of minor bias. Future implementations might leverage the inherent properties of the Markov model itself to estimate latent states and missing observations.^15^ This is particularly important for home monitoring systems, where the quality or continuity of recordings may be impaired, and robust methods for handling missing data are essential for reliable state estimation.

It is important to acknowledge that the assumption of linearity in PCA and ICA may not reflect the full complexity of brain dynamics.^1,35^ Similarly, exploring additional feature sets beyond the spectral domain, such as, non-linear time-domain features, metrics of critical brain dynamics^46,47^ or connectivity measures^48^ could offer deeper insights or highlight different processes. Furthermore, integrating additional modalities, including those derived from metabolic (breathing, temperature, etc.) or hemodynamic signals, may provide a more holistic view of brain-body interactions during sleep and wakefulness.^12,49,50^

The training and validation were performed on a cohort of healthy individuals. While this ensures that the discovered patterns are robust in typical sleep, clinical applications will require extension to diverse populations and conditions. Similarly, reducing the number of electrodes or including different sensor modalities should be used to test the method’s adaptability and robustness.

Applying the method to patient populations with neurological or psychiatric conditions will help validate its clinical utility and sensitivity to pathological changes in sleep. It may reveal particular discrepancies that may be reflected in the feature map or temporal structure of sleep. Integrating these techniques into home-based sleep monitoring systems, wearable devices, or epilepsy monitoring units may facilitate long-term, effortless sleep assessments that can add a prognostic and diagnostic value to these evaluations. This would significantly reduce labeling overhead and empower clinicians and researchers to study sleep over extended periods and large populations without sacrificing data quality or interpretability.

## Conclusion

This study highlights the power of data-driven, unsupervised methods in sleep research. By combining PCA, GHMM, and ICA, we captured and characterized essential features of sleep architecture, replicating and extending conventional staging methods, while revealing new subtleties in temporal dynamics and underlying physiological processes. The robust alignment with conventional scoring, achieved using minimal labeled data, demonstrates the feasibility of scalable, reproducible, and more objective sleep analysis.

In an era where big data and advanced computational tools are reshaping biomedical research, our approach offers a flexible and transparent framework that enriches the understanding of sleep architecture and state transitions. With further refinement, validation, and integration into clinical practice, these methods could improve the diagnosis and management of sleep disorders, advance our understanding of sleep’s role in neurological conditions, and ultimately contribute to better health outcomes.

## Acknowledgments

We would like to acknowledge Prof. Yuval Nir for his advice and suggestions on following this direction.

