## Supplementary material for "The Hypno-PC: Uncovering Sleep Dynamics through Principal Component Analysis and Hidden Markov Modeling of Electrophysiological Signals": Suplemental information

### Supplemental Materials

#### Figure S1 – Principal Component Timeline and Sleep Stages at 4-Second Resolution.

This figure qualitatively illustrates the relationship between the components derived from the 4-second segment decompositions of sleep data. While PC1 still aligns with the hypnogram, the increased temporal resolution introduces more intra-stage variation and weaker stage separations.

**A.** The manually scored hypnogram for the first subject (EPCTL01) is displayed, providing a reference for conventional sleep stage distribution throughout the night.

**B.** PC time series and cross-subject distributions. Left side: Time series of Principal Components 1 through 5 (PC1–PC5) for the first subject, highlighting the substantial similarity between PC1 and the hypnogram values. Right side: The cross-subject empirical distribution for each sleep stage within each component demonstrates the consistency of this representation across individuals. The overall structure is similar to the one found for the 30-second segments, albeit with more within-stage variability.

**C.** Distribution of sleep stages in PC space. The cross-subject representation in the low-dimensional principal PC space shows how different sleep stages occupy distinct regions. However, the separation between Non-Rapid Eye Movement Stage 1 (N1) and Rapid Eye Movement (REM) sleep is not well defined, as evident in panels A and B.

#### Abbreviations:

PC: Principal component, N1–3: Non-rapid eye movement stage 1–3, REM: Rapid eye movement sleep, W: Wake.

#### Figure S2- Principal Component Filters Maps at 4-Second Resolution

This figure illustrates the principal component (PC) filters derived from 4-second segment decompositions of sleep data. PCs 1–3 are nearly identical to the maps obtained from 30-second segments. They similarly relate to different sleep stages, maintaining alignment with the hypnogram and effectively distinguishing major sleep stages. PCs 4 and 5 show similarities; however, the spatial patterns captured by the 4-second windows appear more diffused.

Positive values in PC4 are strongly REM sleep and negatively related to N2, as shown on the 30-second time scale. However, in contrast to the 30-second segments, the positive representation of N1 in this component is distributed around zero and becomes nonsignificant.

In PC5, positive values are strongly related to REM and N1 in most subjects, while W, N2, and N3 are predominantly associated with negative values.

This creates a contrast pattern that inversely resembles PC3, but relies on a different set of features. Several features share a pattern, such as slow waves and delta activity, regardless of this inversion. This highlights the importance of analyzing different frequency bands in the context of changes in other bands. For example, an increase in slow waves combined with increases in alpha, heart rate, muscle movement, and eye movement emphasizes wakefulness, whereas the same increase in slow waves coupled with increases in beta activity and reduced EMG emphasizes REM and N1 stages.

The higher temporal resolution of 4-second segments introduces greater intra-stage variability and weaker separations between sleep stages. This results in more heterogeneous states for N1 and REM sleep, potentially reflecting transient substates that occur within the broader 30-second epochs labeled as Wake (W), REM, N1, and N2.

#### Abbreviations:

PC: Principal component, N1–3: Non-rapid eye movement stage 1–3, REM: Rapid eye movement sleep, W: Wake, sigma: Sigma frequency band, theta: Theta frequency band, alpha: Alpha frequency band, delta: Delta frequency band, gamma: Gamma frequency band, EMG: Electromyography

### Figure S3 – GHMM Cross-Validation Results for 30-Second and 4-Second Segments.

This figure presents the cross-validation results for 30-second (Panel A) and 4-second (Panel B) segments performed to evaluate the optimal number of states in the range of 2–15 hidden states.

**A.** Cross-validation results in the 30-second resolution. This panel shows the trends of the Bayesian Information Criterion (BIC) and Akaike Information Criterion (AIC) across different numbers of hidden states. Both metrics gradually decline as the number of hidden states increases to four states, indicating an improvement in the model fit to the test data. After four states, the BIC measure starts to increase gradually, indicating that the cost of model complexity is not balanced by the fit quality of the model to the data.
Although not utilized for feature selection, the analysis of Kappa and accuracy metrics reveals that four hidden states yield the best median agreement with the conventional sleep labels. This state also shows lower cross-subject variability, indicating a more consistent model. This suggests enhanced reliability in the alignment of the hidden states with the sleep stage labels.

B. Cross-validation results at the 4-second resolution. At this resolution, both BIC and AIC exhibit a gradual decline with increasing hidden states. Both metrics stabilize at seven states, and no significant improvement of the BIC and AIC values is seen with a higher number of states. The median kappa and accuracy metrics are highest with seven states. However, higher states show lower cross-subject variability, which may indicate models that are more stable in terms of alignment with sleep labels. The variability reduction was not evident in the AIC and BIC metrics; thus, this was not considered in the state selection.

#### Abbreviations:

BIC: Bayesian information criterion, AIC: Akaike information criterion, Kappa: Cohen's kappa coefficient, GHMM: Gaussian hidden Markov model.

#### Figure S4 – Transition and Stability Comparison Using GHMM Label Assignments.

This figure extends the example presented in Figure 4 by applying a shared Gaussian hidden Markov model (GHMM) at the 4-second time scale, which consistently incorporates a distinct N1-like state. This approach enables the estimation of state stability and transition probabilities for all conventional sleep stages. We first used the GHMM to assign hidden states to data from subjects EPCTL16 and EPCTL28 and then mapped these hidden-state assignments onto traditional sleep-stage labels. Each label sequence (hidden states vs. sleep stages) was subsequently used to analyze transition probabilities and stage stability, producing two transition plots per subject.

A. GHMM Assigned Sleep Stages Transition Estimation: For subject EPCTL16, we observe reduced stability in Wake (W) and Rapid Eye Movement (REM) sleep, reflected by higher transition probabilities away from these states. This finding aligns with the manual labeling (Figure 3), wherein W and REM also appear to act as weaker attractors. EPCTL16 additionally exhibits increased transitions from REM to N1 and N2, surpassing both the rates of EPCTL28 and the average transition (Figure 5), suggesting that there is more pronounced REM fragmentation in EPCTL16. Although fewer transitions from REM to N2 are noted in the manual labels, the GHMM-based approach more readily captures these events. A minor direct W-to-REM pathway is also observed in EPCTL16, which is not captured by the shared GHMM model (Figure 5). Although it remains sub-diagnostic, it is important to note that the earlier-than-typical REM onset (58.5 minutes) in this subject may indicate heightened REM pressure or narcolepsy-like features in this subject.

B. The GHMM hidden-state transition plots for the same two subjects similarly illustrate reduced REM stability in EPCTL16, with increased transitions from REM into N1 and N2. A salient observation is that the W-to-REM transition appears to originate from a specific high-alertness wake sub-state (hidden state 3 in Figure 5, lower panel) rather than from quiescent wakefulness. The GHMM also reveals transitions between an N3-related sub-state and a REM-related one, suggesting that there are more permeable boundaries around the REM sub-state. This heightened connectivity into and out of REM may signify a subclinical impairment of REM stability in subject EPCTL16. Overall, these findings demonstrate that GHMM-based labeling can capture subtle transition dynamics not readily discerned from a manual hypnogram alone.

##### Abbreviations:

GHMM: Gaussian hidden Markov model, REM: Rapid eye movement sleep, N1–3: Non-rapid eye movement sleep 1–3, W: Wake

#### Supporting Results: Identifying Sources Using Independent Component Analysis (ICA)

To further dissect the low-dimensional representation obtained from principal component analysis (PCA), we applied ICA to the five PCs. Unlike PCA, which prioritizes orthogonal directions of maximal variance, ICA seeks to isolate statistically independent signal sources,[1] and is often employed at the raw electroencephalographic (EEG) level to separate muscle activity, electrocardiography (ECG) components, electrooculography (EOG), and line noise.[2] In this work, by unmixing the PCs, ICA uncovers subtle influences that may overlap in PCA space, providing additional granularity in interpreting the temporal and spatial organization of sleep stages, which has been suggested as highly important in the development of sleep research.[3] Figures S5 and S6 illustrate the time courses and filter maps of the five independent components (ICs) derived through ICA. While the PCA findings align strongly with recognized sleep-stage categorizations, ICA enriches this perspective by revealing components that may not strictly adhere to orthogonal constraints. This approach refines our understanding of how partially independent physiological processes (e.g., arousals, phasic REM events, muscle surges) contribute to the multifaceted nature of sleep.

#### Figure S5 – Independent Component Timeline and Sleep Stages at 30-Second Resolution

The time series of the five ICs derived from ICA are shown for a representative subject, compared to the manual hypnogram presented on the top panel.

**IC1:** Mirrors in the hypnogram capturing both general trends and short awakening events, with lower variability within N2 and N3 compared to PC1 (Figure 1), indicating a more stable representation of the core sleep stages.

**IC2:** Spans a gradient from N3 (negative) to N1/N2 and REM sleep (positive), reflecting the interplay between slow-wave sleep and more activated states. It exhibits greater variability in the latter part of the night, which may represent changing network activity in these stages.

**IC3:** Primarily differentiates wakefulness from N3. It shows the highest variability in wakefulness, even though brief awakenings are not strongly captured, implying a quiet, low-movement sub-state of wakefulness. Like IC2, we see greater variability in the lighter sleep stages and wakefulness compared to N3, reflecting activity fluctuations within these stages.

**IC4:** Emphasizes short awakenings and muscle surges (Figure S6). All sleep stages fall near small negative values, with only a small subset of segments mapping strongly into positive values, which may reflect brief arousals.

**IC5:** Wakefulness is mainly contrasted with certain aspects of REM, exhibiting high variability in both states. This may reflect network activity related to phasic/tonic REM[4], dream experiences[5], or transitions in and out of wakefulness.[6]

**Lower panel:** The left plot displays ICs 1–3 in a low-dimensional space, and the right plot displays ICs 1, 4, and 5. In both plots, data points are confined to a manifold-like subspace. Color-coding by sleep labels reveals a gradual spatial representation of the stages within this manifold.

#### Abbreviations:

ICs: independent components; PC: principal component; N1-3: Non-rapid eye movement 1-3, REM: rapid eye movement

### Figure S6 - Independent Component Filters Maps at 30-Second Resolution

Topographic feature-weight maps for each IC illustrate how EEG electrode locations, frequency bands, and non-EEG measures (EOG, EMG, ECG) contribute to these independent processes.

**IC1 Map:** High positive weights for posterior alpha, fronto-central theta, and high-frequency EEG signals typical of wakefulness. Negative weights for central posterior sigma and slow waves are associated with N2/N3. Compared to PC1, IC1 shows a more specific spatial distribution, implying finer spatial feature discrimination.

**IC2 Map:** The positive direction indicates N1, N2, and REM, with increased spectral slope, entropy, and a blend of delta, theta, and high sigma. The negative direction emphasizes frontal slow waves (N3) and posterior alpha (wake), placing wakefulness in a more neutral or modestly negative position. EOG power is associated with the negative direction; however, given the limitations in our analysis, it is hard to distinguish this trend from the increase in frontal slow wave power.

**IC3 Map:** Contrasts global alpha-sigma (wake) with delta/theta waves (REM/N3). Exhibits minimal correlation with EOG, EMG, or ECG changes, indicating that it primarily captures specific EEG modulations. This component indicates high variability in W, REM, and N2 with global (though more posterior) shifts in 1–4 Hz and 8–15 Hz frequencies.

**IC4 Map:** Negative values are associated modestly with stable sleep states (N2, N3), while positive values characterize wakefulness. Elevated EMG, EOG, heart rate, high sigma, and lateral gamma activity link IC4 to short arousals and active wakefulness.[3,7,6]

**IC5 Map:** Separates wakefulness and REM through frontal gamma, alpha, posterior power, and an increased muscle/heart rate in the wakefulness direction, contrasted with frontal slow waves, entropy, posterior beta/sigma, and an increase in lateral eye movements in the REM direction. Negative values may capture pontine-genicular-occipital (PGO) waves tied to phasic REM.[5,8]

These IC-derived time courses and filter maps emphasize the complex interplay of cortical activation patterns, muscle surges, and autonomic fluctuations.[9,10] Each independent component isolates partially independent processes—ranging from robust wake-to-REM differentials to subtler sub-states of N3 or transient arousal events—enhancing our grasp of the latent networks underpinning sleep continuity and disruption.[3,7] While ICA does not significantly revise the boundaries of sleep stages identified by PCA and GHMM, it informs finer distinctions, highlighting the complexity of nocturnal brain and body states. Ultimately, the ICA findings contribute an additional layer of detail to our comprehensive portrait of sleep's spatiotemporal organization.
