## Supplementary figures and images for "The Hypno-PC: Uncovering Sleep Dynamics through Principal Component Analysis and Hidden Markov Modeling of Electrophysiological Signals"

### Figure S1

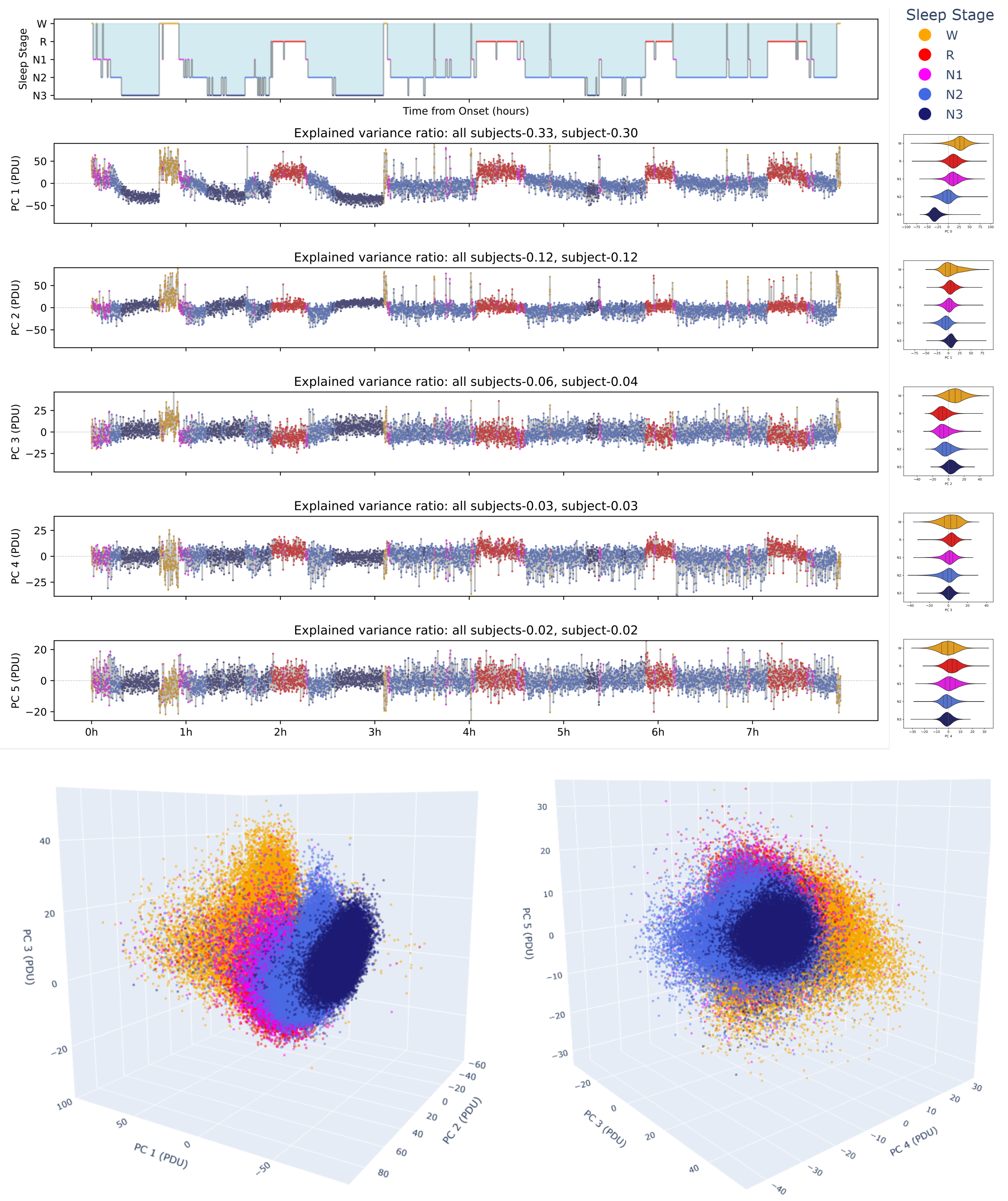

### Figure S2

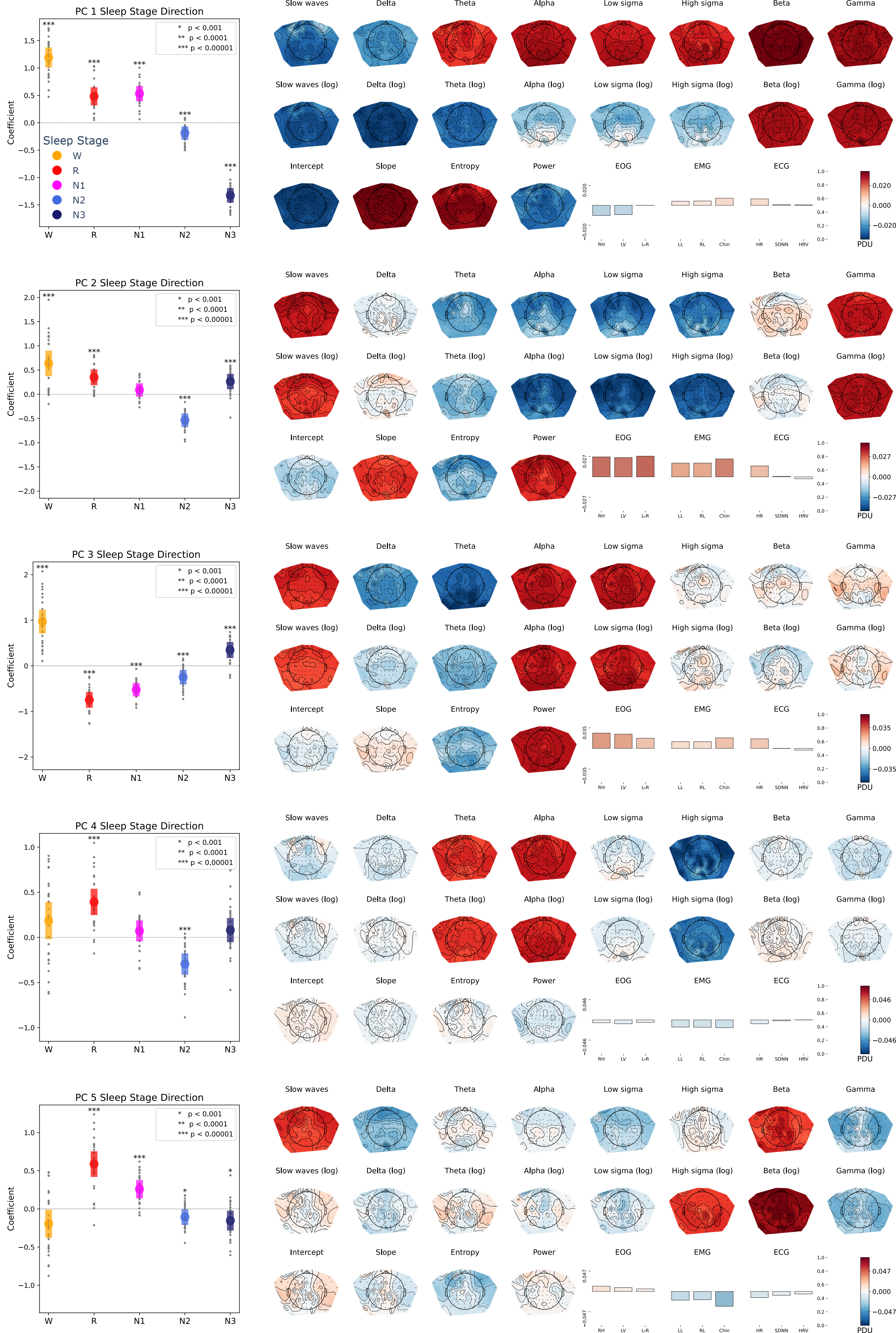

### Figure S3

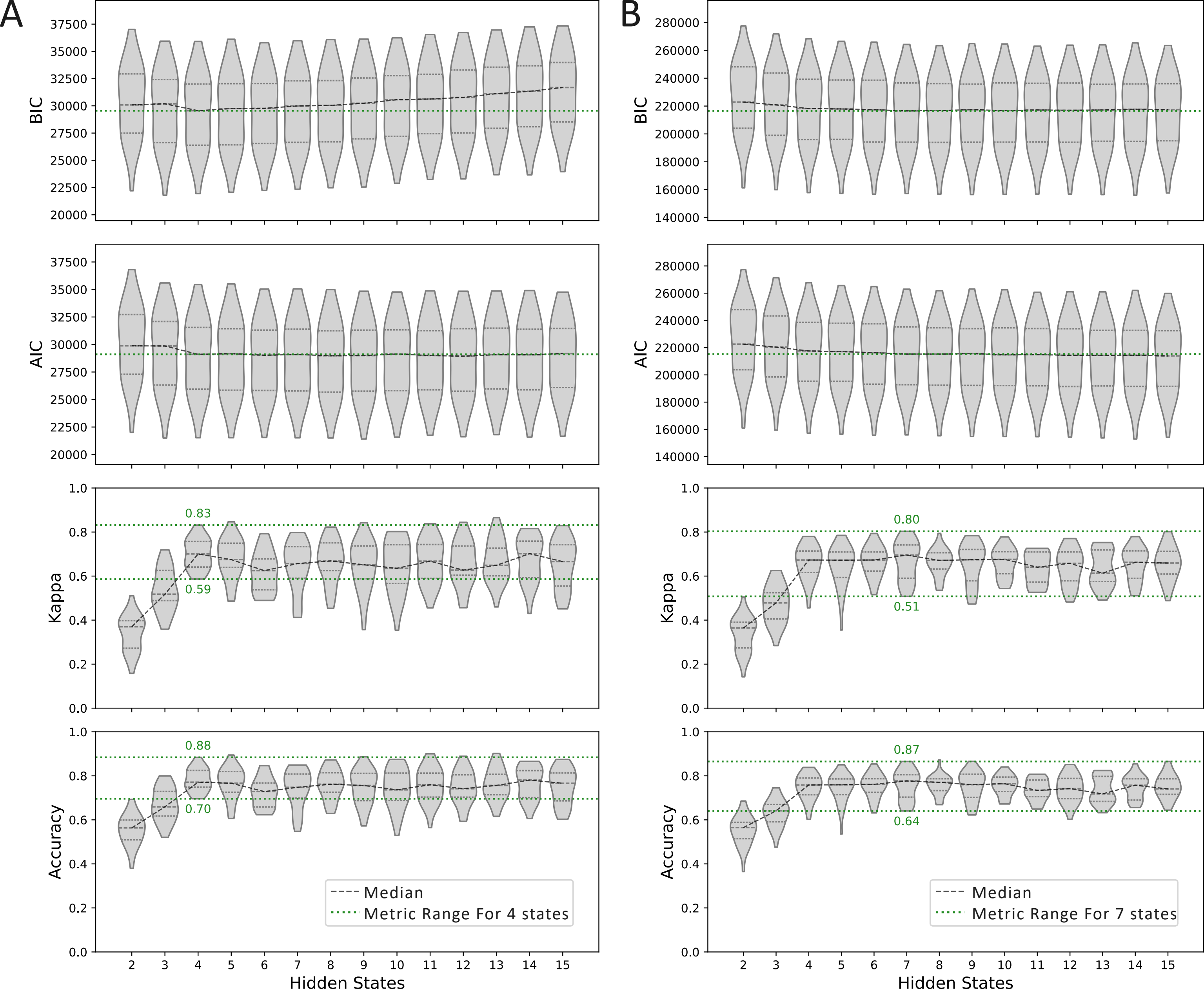

### Figure S4

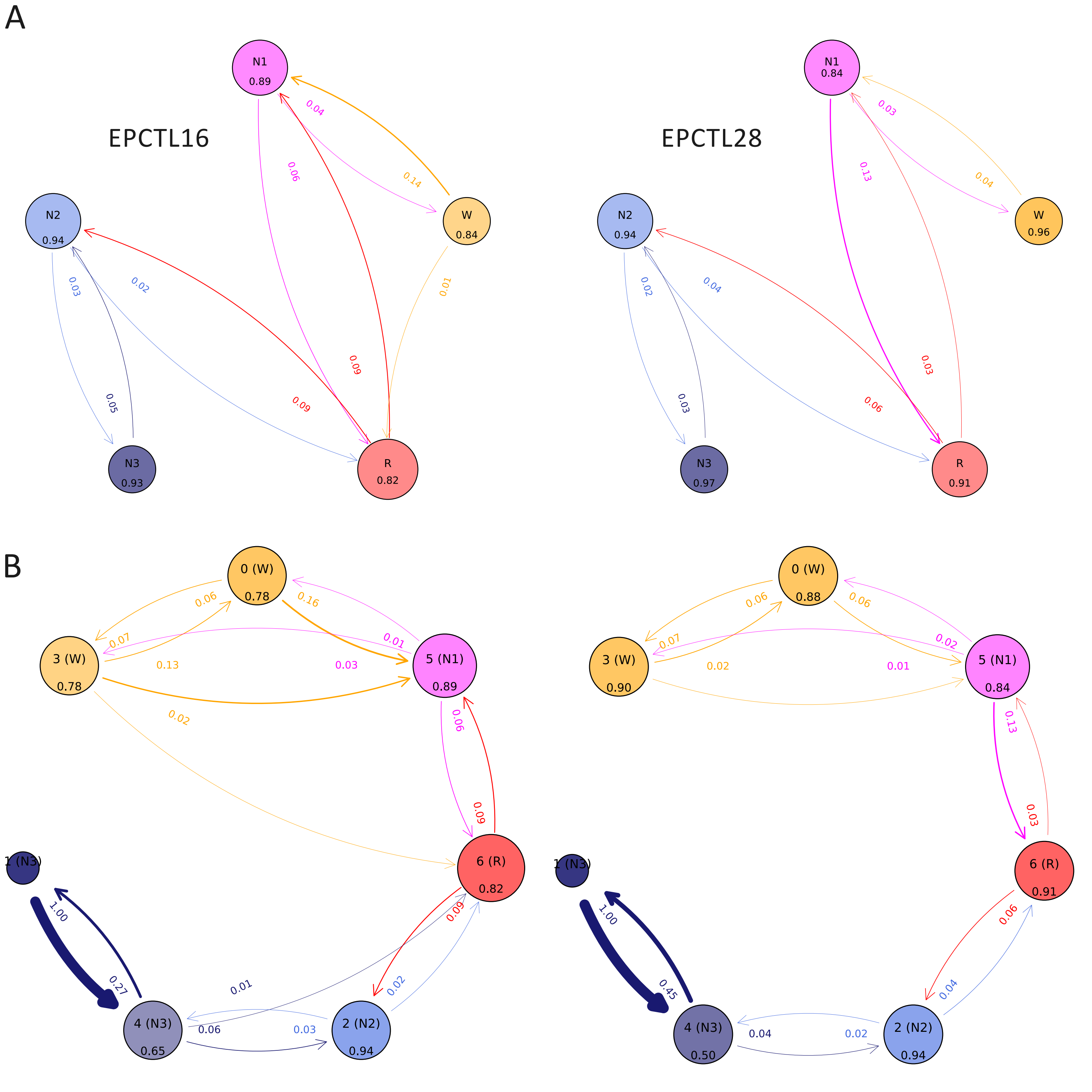

### Figure S5

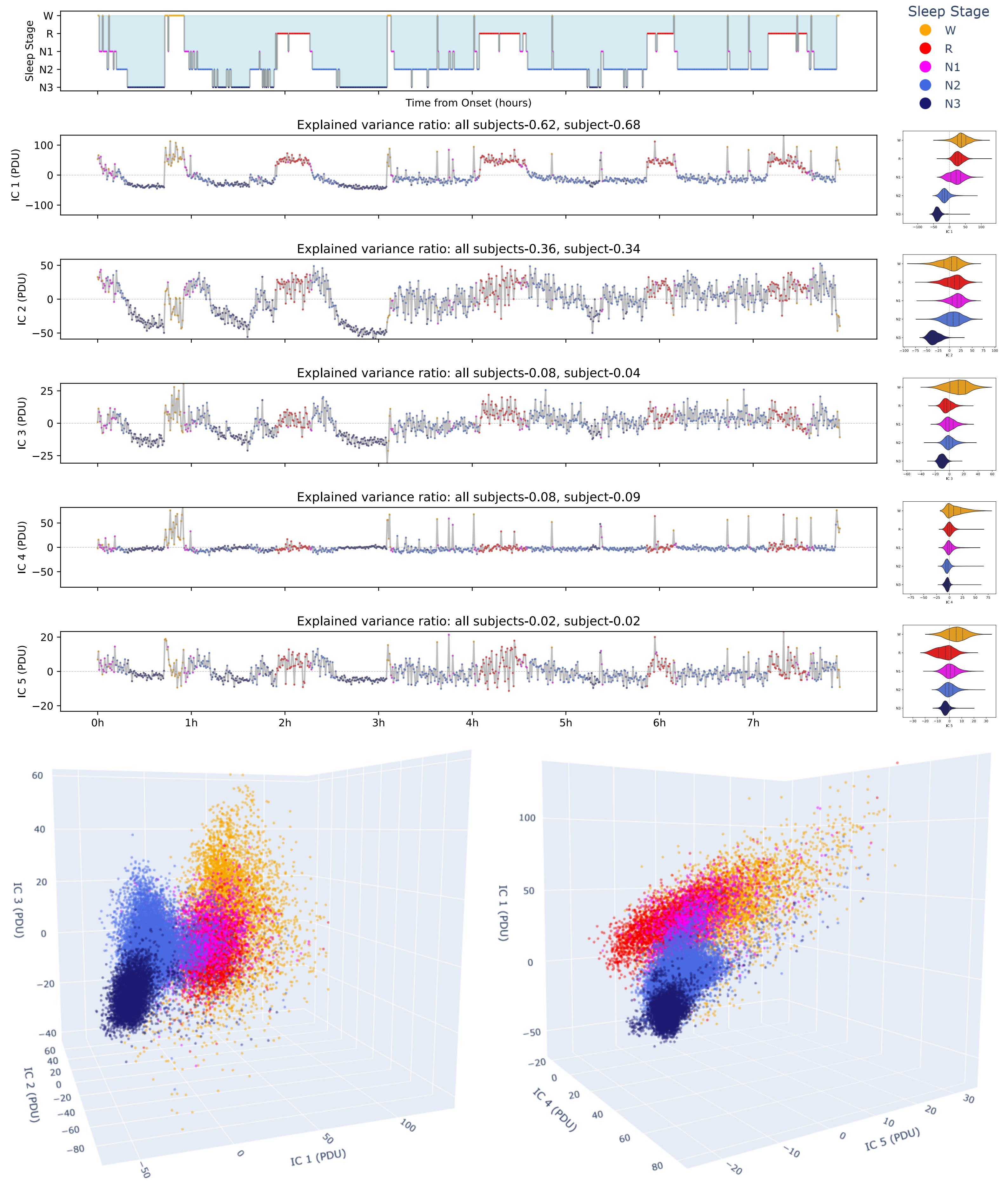

### Figure S6

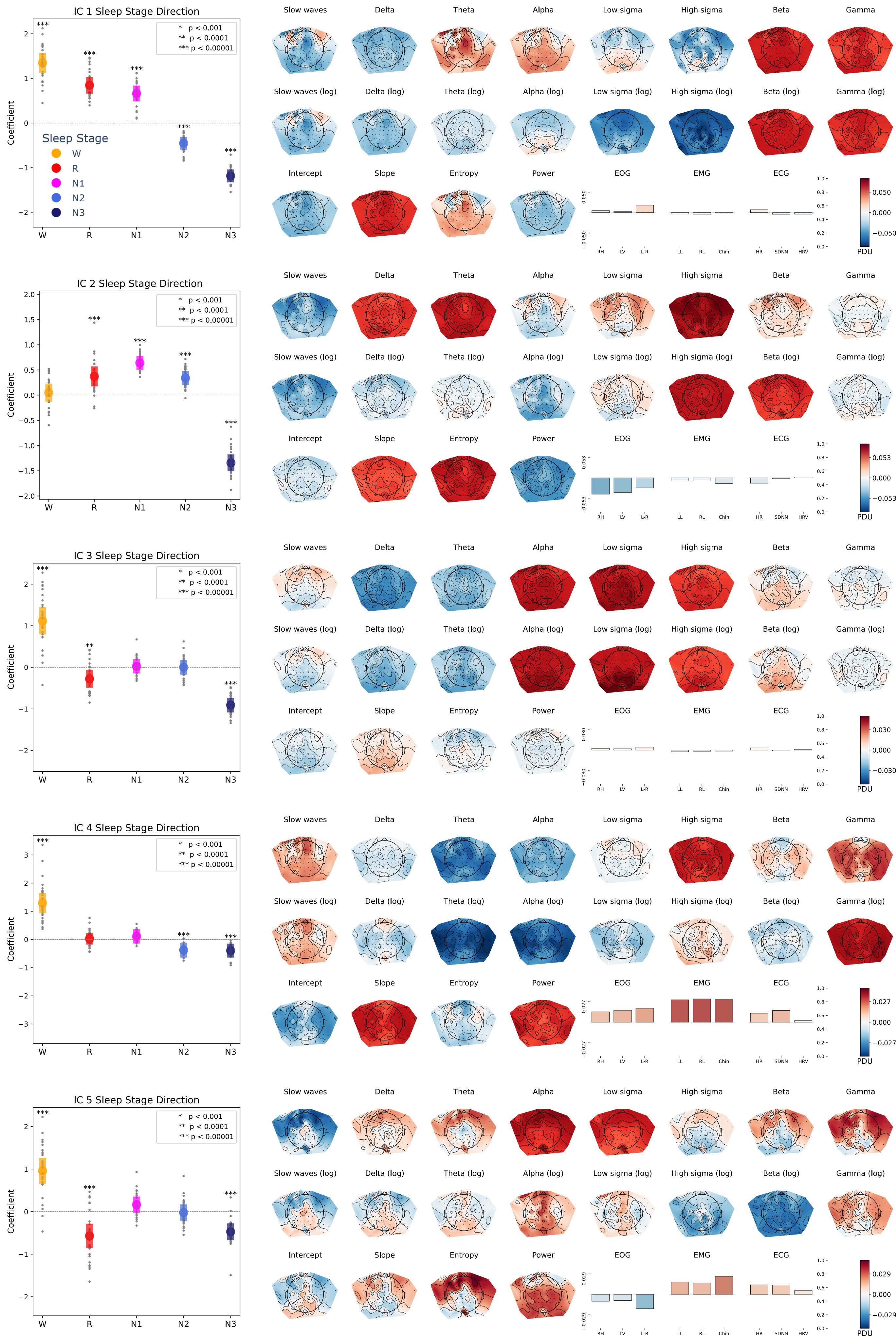
